# Refining the evolutionary time machine: an assessment of whole genome amplification using single historical *Daphnia* eggs

**DOI:** 10.1101/2021.04.19.440325

**Authors:** Christopher James O’Grady, Vignesh Dhandapani, John K. Colbourne, Dagmar Frisch

## Abstract

Whole genome sequencing is instrumental for the study of genome variation in natural populations, delivering important knowledge on genomic modifications and potential targets of natural selection at the population level. Large dormant eggbanks of aquatic invertebrates such as the keystone herbivore *Daphnia*, a microcrustacean widespread in freshwater ecosystems, provide detailed sedimentary archives to study genomic processes over centuries. To overcome the problem of limited DNA amounts in single *Daphnia* dormant eggs, we developed an optimised workflow for whole genome amplification (WGA), yielding sufficient amounts of DNA for downstream whole genome sequencing of individual historical eggs, including polyploid lineages. We compare two WGA kits, applied to recently produced *Daphnia magna* dormant eggs from laboratory cultures, and to historical dormant eggs of *Daphnia pulicaria* collected from Arctic lake sediment between 10y and 300y old. Resulting genome coverage breadth in most samples was ~70%, including those from >100y old isolates. Sequence read distribution was highly correlated among samples amplified with the same kit, but less correlated between kits. Despite this, a high percentage of genomic positions with SNPs in one or more samples (maximum of 74% between kits, and 97% within kits) were recovered at a depth required for genotyping. As a by-product of sequencing we obtained 100% coverage of the mitochondrial genomes even from the oldest isolates (~300y). The mtDNA provides an additional source for evolutionary studies of these populations. We provide an optimised workflow for WGA followed by whole genome sequencing including steps to minimise exogenous DNA.

## Introduction

With recent declines in sequencing costs due to development of high-throughput technologies, whole genome sequencing (WGS) has emerged as an important molecular tool in evolutionary biology, and has been applied to a plethora of different biological systems (Dettman et al., 2012; Ellegren, 2014; Hohenlohe, Hand, Andrews, & Luikart, 2018; Stiller & Zhang, 2019). WGS allows the analysis of genetic variation at thousands of genomic loci to test relationships between phenotypic and genotypic adaptations in genome-wide association studies (GWAS) (De La Torre, Wilhite, & Neale, 2019; Rajpurohit et al., 2018; Sella & Barton, 2019). At the population level, and in particular if long-term time series data including from ancient DNA are available, WGS can provide invaluable genomic detail, shedding light on evolutionary patterns and processes (Leonardi et al., 2017; Parks et al., 2015).

Understanding how individuals and populations adapt to their environment is one of the most compelling and challenging tasks in evolutionary ecology, especially regarding the current unprecedented environmental change. A unique approach gaining momentum is the study of propagules of various plant or animal taxa preserved in layered aquatic sediments (Ellegaard et al., 2020; Orsini et al., 2013). These archives, consisting of dormant stages (e.g. eggs, seeds, cysts) with DNA degraded to varying degrees, as well as of hatchable propagules with intact DNA allow the direct observation of evolutionary change across centuries or even millennia (Brede et al., 2009; Cordellier, Wojewodzic, Wessels, Kuster, & von Elert, 2021; Frisch et al., 2014; Härnström, Ellegaard, Andersen, & Godhe, 2011; Mergeay, Verschuren, & De Meester, 2006; Pollard, Colbourne, & Keller, 2003; Weider, Lampert, Wessels, Colbourne, & Limburg, 1997), potentially at genomic resolution of individual isolates. The exploitation of such resources together with modern molecular tools is instrumental in the study of evolutionary processes over thousands of generations in relation to environmental change.

As one of the notable examples, the population genetics of the ecological and genomic model *Daphnia* (Crustacea, Cladocera) has been studied over historical time frames and associated with changes in the lake environment, genotyping either individual eggs (Limburg & Weider, 2002; Brede et al., 2009; Orsini, Spanier, & De Meester, 2012; Frisch et al., 2014, 2016) or by whole genome sequencing of pooled egg DNA (Cordellier et al., 2021). Whole genome sequencing (WGS) of individual eggs would allow population genomic studies at high-resolution, however, success of WGS of individual dormant eggs so far has been limited (Lack, Weider, & Jeyasingh, 2018).

A major obstacle for WGS of individual dormant *Daphnia* eggs is their minute amount of DNA. These eggs contain an embryo in the late blastula stage (Chen et al, 2018; von Baldass, 1941) given an estimated haploid genome size of ~200 Mb (Colbourne et al., 2011) and around 2000 cells in a dormant embryo of *D. pulex* (von Baldass, 1941), the DNA content can be estimated at ~800 pg in a diploid embryo. The situation is exacerbated for historical dormant *Daphnia* eggs or those of other taxa, due to DNA degradation, posing additional problems for DNA sequencing (Rizzi, Lari, Gigli, De Bellis, & Caramelli, 2012). Although it is possible to combine eggs from individual sediment strata for a pooled sequencing approach, such a strategy leads to information loss on individual genotypes and accuracy of population genomic parameters such as F_ST_ estimates (Dorant et al., 2019). An alternative approach to gain sufficient amounts of genetic starting material is by performing whole genome amplification (WGA). This method uses cell material without prior DNA extraction, thus minimising potential loss of DNA during the extraction process and amplifies genomic DNA from extremely low starting concentrations in the picogram range. However, a previous study that applied WGA to individual, dormant *Daphnia* eggs had limited success with only one of three eggs producing amplified *Daphnia* DNA (Lack et al., 2018).

Multiple displacement amplification (MDA), a widely used PCR-free WGA method, utilizes a high fidelity φ29 DNA polymerase which extends from hexamer primers that randomly bind to targets across the genomic template (Dean et al., 2002; Dean, Nelson, Giesler, & Lasken, 2001). This results in the generation of large DNA products, with an average length of ~10kb (capable of reaching over 100 kb), that can have a strong coverage of the target genome (Blanco et al., 1989; Handyside et al., 2004; Lasken & Egholm, 2003; Paez, 2004). MDA is often favoured over PCR-WGA techniques, for example degenerate oligonucleotide-primed PCR, as PCR-based methods can result in the production of small DNA fragments (>1 kb) (Telenius et al., 1992; Wells, Sherlock, Handyside, & Delhanty, 1999; L. Zhang et al., 1992) that contain a number of non-specific amplification artifacts (Cheung & Nelson, 1996). Additionally, PCR-WGA methods can show a significant amplification bias towards specific loci, and consequently products may not give a complete coverage of loci (Dean et al., 2002). MDA is highly sensitive and is particularly vulnerable to DNA contamination, which can compete or co-amplify with the desired DNA template during WGA and cause issues during downstream analyses (Blainey & Quake, 2011; Woyke et al., 2011). Great care must therefore be taken to eliminate sources of contamination during the amplification step.

Extending a study for whole genome amplification from individual *Daphnia* (Lack et al., 2018), our goal was to develop an optimized WGA-WGS workflow for historical dormant egg isolates including improved decontamination steps. To achieve this, we use recently produced *Daphnia magna* dormant eggs from laboratory cultures (days old) and *Daphnia pulicaria* dormant eggs isolated from lake sediment (between 10-300 years old), and compared two commercially available single cell WGA kits based on MDA technology. We test the success of reducing exogenous DNA through the application of different concentrations and durations of bleach, or several washes with PBS to samples prior to WGA.

In a sequencing experiment, we analyse mapping efficiency and genome-wide read distribution and compare these between species and eggs of various age for a total number of 16 dormant eggs aged up to 300 years old. We compare read distribution patterns, coverage breadth and uniformity, and identify contaminants. Finally, we test the utility of these kits for detecting genomic variants in both nuclear and mitochondrial genomes.

## Materials and Methods

### Egg collection

All eggs used for whole genome amplification were isolated from ephippia of two species: *Daphnia magna* Straus, 1820, and *Daphnia pulicaria* Forbes, 1893. For *D. magna*, we used eggs from recently produced ephippia that are routinely removed from laboratory cultures maintained in the *Daphnia* facility of the University of Birmingham, UK (DM1-DM7, unknown origin). Ephippia of Arctic, triploid populations of *Daphnia pulicaria* were collected in 2015 from sediment of two lakes in West Greenland (Kangerlussuaq area). Details on the lakes and sediment dating can be found in Dane, Anderson, Osburn, Colbourne, & Frisch(2020). Briefly, we sampled ephippia from sediment corresponding to several historical time periods in two lakes: Lake SS4 (Braya Sø): ca. 2010, ca. 1880, ca. 1720, and Lake SS381: ca. 2010, ca. 1840 (Table 1).

**Table 1.**
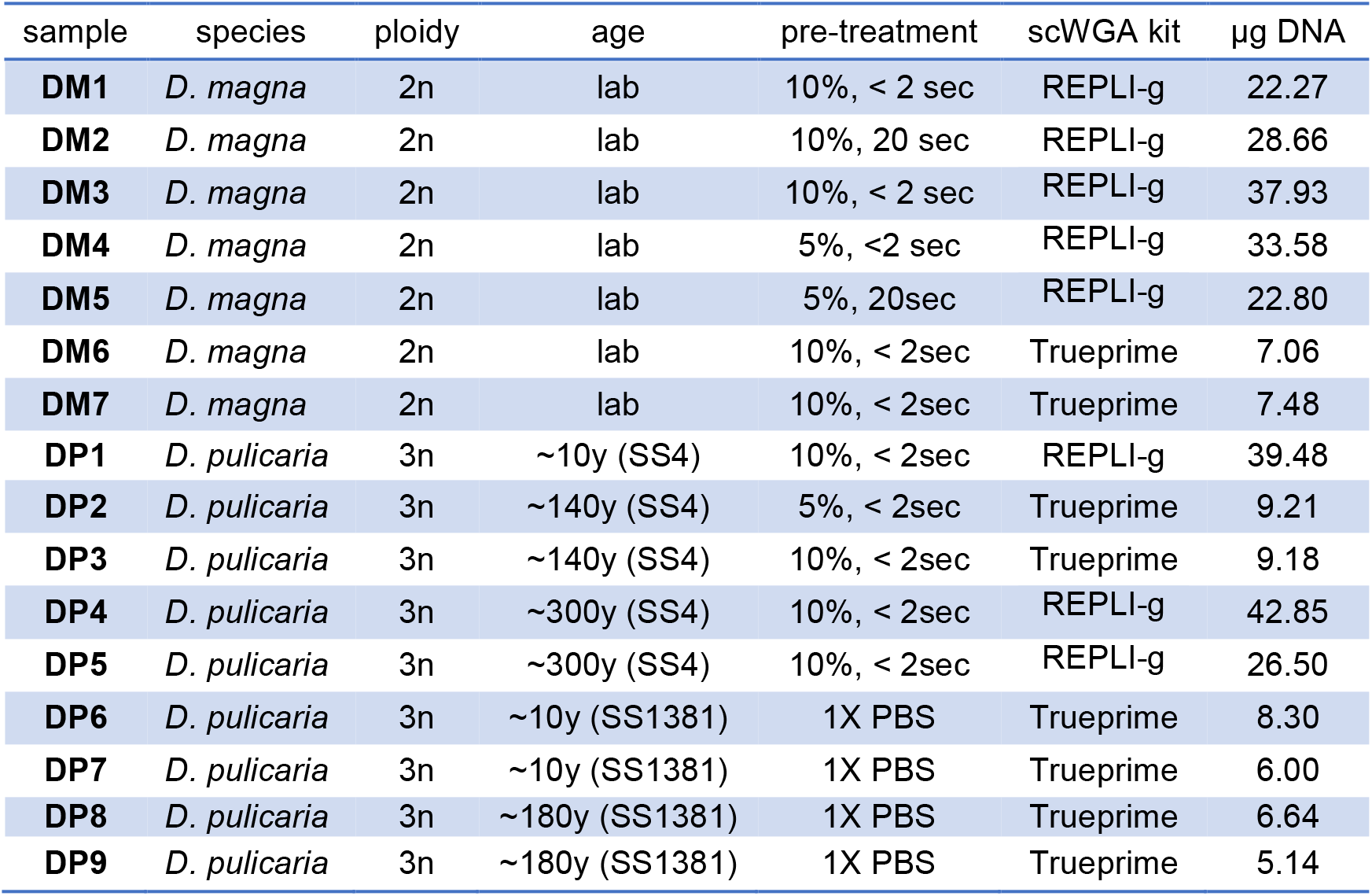
Species identity and specifics of dormant eggs used as template in whole genome amplification. Additional information is given on pre-treatment (% of the bleach solution and exposure time, or 1X PBS buffer, e.g. “10% <2 sec” indicates exposure for less than 2 seconds to a 10% bleach solution made from 12% industrial bleach), the applied WGA kit and the total product of WGA-DNA obtained. SS4 and SS1831 are two lakes in West Greenland near Kangerlussuaq (for details see Methods) from which sediment cores with ephippia were extracted.

### Pre-WGA preparation and cleaning of Daphnia eggs

Eggs were removed from ephippia (decapsulated) and transferred to sterile 1X PBS shortly before use. Decapsulated eggs were inspected under a stereomicroscope to ensure that eggs were in good condition (judged by colour and appearance). Visually undamaged eggs were washed in a 5% or 10% wash solution made from industrial strength (12%) bleach. Exposure to the wash solution was either instantly (<2 seconds) or for 20 seconds (Table 1), followed by five separate rinses in a sterile 1X PBS buffer to remove any remaining bleach. Alternatively, eggs were washed by five to eight rinses in 1X PBS, by placing a row of PBS droplets on a glass slide, and washing each egg individually by carefully and repeatedly drawing them up with a pipette (sterile tip) in each of the droplets. Rinse controls contained the PBS solution used for the last rinse, while negative controls contained sterile PBS. Bleaching (including the final PBS rinse) was performed in a SCANLAF Mars Safety Class 2 laminar flowhood in sterile conditions. The alternative procedure of PBS rinsing was performed in a clean, dedicated room with thorough bleaching of all surfaces prior to processing eggs. Bleached and rinsed eggs were kept on ice for brief periods in sterile 1X PBS until further processing.

### Whole Genome Amplification

Whole Genome Amplification (WGA) was performed using two PCR-free kits: Expedeon TruePrime™ Single Cell WGA kit (from hereon: TruePrime), and Qiagen REPLI-g Single Cell Kit (from hereon: REPLI-g). Positive controls (extracted *Daphnia* DNA) were included in WGA. Rinse controls and negative controls were included to monitor possible amplification of contaminating DNA. Prior to WGA, egg membranes were pierced with a sterile 10 μl pipette tip to allow exposure of embryonic cells, and kept in the respective amount of 1X PBS buffer required for the first step of the reaction in each test kit. DNA concentration in WGA products were quantified with a microplate Reader (Tecan infinite F200 pro), or a Qubit^®^ 2.0 Fluorometer (Invitrogen) and the Qubit^®^ dsDNA HS Assay kit (Invitrogen). The size distribution of WGA products was determined by agarose gel electrophoresis to analyse the impact of pre-treatment steps (bleaching or washing in PBS) on the DNA fragments produced by WGA.

### Whole genome library preparation and sequencing

WGA samples DM1 - DM7 (*Daphnia magna* eggs) and DP1 - DP5 (*Daphnia pulicaria* eggs) were used to prepare single-end (SE) libraries with an insert size of 300 bp using a PCR-free workflow with the KAPA HyperPrep Kit (Roche) following manufacturer’s instructions. Sequencing of 100 bp SE libraries was performed on the Illumina HiSeq2500 platform at the Environmental Omics sequencing facility, University of Birmingham, UK. WGA samples DP6 – DP9 (*Daphnia pulicaria* eggs) were used to prepare paired-end (PE) libraries with an insert size of 350 bp with the TruSeq DNA PCR-free gel-free library preparation kit (Illumina) according to the manufacturer’s instructions. Paired-end library preparation and sequencing (150 bp PE libraries) was performed at Edinburgh Genomics, The University of Edinburgh, UK.

### Bioinformatic and statistical analysis

All analyses involving R packages were completed with R version 3.6.2 (R CoreTeam, 2019).

*Quality control and mapping.* We used FastQC (Andrews, 2015) to check read quality, followed by adapter trimming and removal of leading and trailing low quality bases with Trimmomatic (Bolger, Lohse, & Usadel, 2014). Following quality control, reads were mapped using BWA-mem with default settings (Li & Durbin, 2009) to the respective reference genome assembly (*Daphnia magna* genome assembly daphmag2.4, GenBank accession GCA_001632505.1; *Daphnia pulex* genome assembly (http://genome.jgi.doe.gov/Dappu1/Dappu1.download.html) (Colbourne et al., 2011), *Daphnia pulex* mitochondrial genome, Genbank Accession NC_000844 (Crease, 1999). Mapping statistics were computed with Qualimap (García-Alcalde et al., 2012) prior to variant calling. Duplicate reads were removed from mapped reads using MarkDuplicates from the Picard Toolkit (Broad Institute, 2019).

*Nuclear DNA variant calling and analysis.* Nuclear Single Nucleotide Polymorphisms (SNPs) were called in *Daphnia magna* using the available SE libraries. SNPs in the triploid Arctic *D. pulicaria* eggs were called only from PE samples because sequencing depth of SE samples was insufficient for calling variants in a triploid organism (Maruki & Lynch, 2017). Nuclear and mitochondrial variants were called with freebayes v. 1.3.2 (Garrison & Marth, 2012), excluding reads with a mapping quality < 40, base quality < 24 and a minimum alternate allele fraction of 0.01, ploidy = 2 (*D. magna* ncDNA, *D. pulicaria* mtDNA) and ploidy = 3 in *D. pulicaria* ncDNA. After variant calling, nuclear SNPs were hard-filtered with vcffilter (Garrison, 2016) applying all of the following settings: “QUAL > 1” to ensure the exclusion of variants with very low quality, “QUAL / AO > 10” to include only variants where each observation contributes at least 10 log units (~Q10 per read), “SAF > 0 & SAR > 0” to avoid strand bias, and “RPR > 1 & RPL > 1” to require at least two reads on each side of the variant.

Genomic positions with high-confidence SNPs present in one or more samples were compared between selected samples to assess the percentage of loci that could be called in all selected samples (i.e. that were amplified and sequenced at the depth required for genotyping). This comparison was used primarily to estimate the repeatability of WGA and subsequent WGS, and thus for the resulting capacity to call variants at multisample level. For *D. magna*, we compared the four samples with the highest number of SNPs (two REPLI-g amplified samples: DM2, DM3, two TruePrime amplified samples: DM6, DM7). For *D. pulicaria*, we compared all four PE samples (only TruePrime amplified). Results were visualised with the R packages eulerr v. 6.1.0 (Larsson, 2020) and ggVennDiagram v. 0.3 (Gao & Yi, 2019).

Transition-to-transversion ratios for SNPs (Ti:Tv) were calculated after applying a minor allele frequency (MAF) threshold of 0.05, and not allowing missing data (R packages SeqArray 1.26.2 (Zheng et al., 2017) and SeqVarTools 1.24.1 (Gogarten et al. 2020)).

*mtDNA variant calling and analysis.* To analyse mtDNA, we used all available *Daphnia pulicaria* samples (DP1-DF10). For mitochondrial SNPs, the same filters as for nuclear SNPs were applied except “QUAL / AO > 10” to avoid filtering calls of the alternate allele from samples with paired-end sequencing due to their consistently higher depth compared to the single-end samples. Identity-by-State (IBS) was calculated by applying a MAF threshold of 0.05, not allowing missing data (R package SeqArray 1.26.2 (Zheng et al., 2017)). SNPs in *Daphnia pulicaria* mtDNA were visualised with the packages Circlize v.0.4.11 (Gu et al. 2014), and SNPRelate (Zheng et al., 2012).

*Read distribution.* This analysis focused on reads mapped to the N50 scaffolds of the *D. magna* and *D. pulex* genomes. Coverage depth was normalised between samples of binned reads (bin size 10 kb or 100 kb, normalised reads = number of reads per bin / average number of reads across bins. Normalised read coverage was visualised with the packages Circlize v.0.4.11 (Gu, 2014), and ggplot2 v.3.3.2 (Wickham, 2016). Read distribution was compared within species between samples by correlation analysis (Pearson’s correlation coefficient) of normalised binned reads. For this purpose, we removed a single outlier present in all *D. pulicaria* samples (position 420,001-430,000, scaffold 38). We tested uniformity of read distribution according to the standard model for random sequencing by fitting the distribution of normalised read coverage to a Poisson distribution (Lander & Waterman, 1988).

*Outlier identification and* exogenous *DNA in DF7 and DF8.* Outlier regions were identified by a mapping rate 10 times higher than the mean normalized count. Sequences belonging to these outlier regions were extracted and searched against the nucleotide database with BLASTN. Exogenous DNA was identified in two samples with low mapping efficiency (DP4, DP5). For this, unmapped reads were called from bam files with samtools v1.4 (Li & Durbin, 2009) and converted to paired-end fastq files using bedtools v.2 (Quinlan & Hall, 2010). Following this step, SOAPdenovo-127mer v.2 (Luo et al., 2012) was used to denovo-assemble the unmapped reads with kmer size of 23 bp and default parameters. The resulting contigs were searched against the NCBI nucleotide database with BLASTN (using the command line “*blastn -task megablast -db NCBI_nt_db -query infile -evalue 1e-100 -out outfile -max_target_seqs 1 -num_threads 10 -outfmt “6 qseqid sseqid sciname qlen slen qstart qend sstart send length evalue pident nident mismatch gaps*“). For graphical representation we used Krona (Ondov, Bergman, & Phillippy, 2011).

## Results

### Whole genome amplification

WGA products were separated by agarose gel electrophoresis to determine the impact of pre-treatment steps (bleaching or washing in PBS) on WGA product fragment sizes. Regardless of the pre-treatment process, strong bands for DNA fragments > 10 kb were detected in all samples, suggesting a minimal impact of pre-treatment on WGA product size (examples in Fig. S1). However, WGA products obtained from one of the rinse controls (1X PBS, DNA amplified from the last PBS wash of an unbleached sample) also produced high intensity bands (from 500 bp to >10 kb), which likely resulted from amplified residual DNA carried over from prior washes. WGA of rinse controls obtained from the PBS after bleach-washing did not yield any product, suggesting that the application of bleach completely removes external, exogenous DNA. No amplified DNA was present in any of the negative controls (sterile PBS).

Mean WGA-DNA concentration was lower in the samples amplified by TruePrime (7.38 μg) in comparison with REPLI-g (31.76 μg) (Table 1). These values were within the ranges suggested by the respective WGA kit manufacturers (~40 μg WGA-DNA for REPLI-g, and 3-4 μg when starting from a single cell for TruePrime).

### Read mapping to nuclear and mitochondrial reference genomes

Both single-end (SE) (DP1-DP5) and paired-end (PE) (DP6-DP9) sequencing was performed for *D. pulicaria*, the latter with about 10-fold higher read numbers and related higher coverage depth (Table 2). We did not find a clear pattern of mapping efficiency to the nuclear genome related to the applied pre-treatment in the several-days-old dormant eggs produced in cultures (DM1-DM7). However, in ephippial eggs isolated from sediment, treatment with any of the tested bleach concentrations (DP1 – DP5) was associated with lower mapping efficiency and/or a smaller fraction of the genome covered, suggesting that bleaching might damage egg DNA. This appeared to be independent of the amplification kit, for example in the TruePrime-amplified samples DP2 and DP3 (bleached, ~50% coverage breadth) compared with the similarly aged DP8 and DP9 (not bleached, ~70% coverage breadth, Table 2a).

**Table 2.** Mapping details and number of variants called in a) nuclear DNA of *D. magna* (DM), and *D. pulicaria* (DP), b) mitochondrial DNA of *D. pulicaria.* age = see Table 1; kit = WGA kit (R-g = REPLI-g, TP = TruePrime); Seq = Sequencing strategy (SE = single end, PE = paired end); Number of reads = total number of reads used for mapping; mapped reads= fraction of reads mapped to the respective reference genome; fraction covered = coverage breadth as the percentage of reference genome with at least 1x coverage; average coverage depth = mean number of reads per genomic position; SNP loci = number of single nucleotide variants compared to the respective reference genome.

**a)**
| Nuclear genome *D. magna* and *D. pulicaria* |  |  |  |  |  |  |  |  |
| --- | --- | --- | --- | --- | --- | --- | --- | --- |
| sample | age | kit | Seq | number of reads | mapped reads % | coverage breadth % | coverage depth | SNP loci |
| DM1 | lab | R-g | SE | 8,488,215 | 98.72 | 74.95 | 6 | 301,779 |
| DM2 | lab | R-g | SE | 17,742,882 | 98.76 | 77.43 | 13 | 414,782 |
| DM3 | lab | R-g | SE | 24,069,767 | 98.74 | 78.63 | 18 | 430,909 |
| DM4 | lab | R-g | SE | 38,765 | 55.73 | 1.47 | 0.02 | NA |
| DM5 | lab | R-g | SE | 10,167,137 | 98.64 | 76.36 | 7 | 339,023 |
| DM6 | lab | TP | SE | 6,392,740 | 99.18 | 66.75 | 5 | 99,927 |
| DM7 | lab | TP | SE | 14,800,247 | 99.04 | 78.05 | 11 | 357,847 |
| DP1 | ~10y | R-g | SE | 12,990,625 | 48.36 | 41.45 | 3 | NA* |
| DP2 | ~140y | TP | SE | 12,088,977 | 87.36 | 50.11 | 5 | NA* |
| DP3 | ~140y | TP | SE | 16,643,733 | 84.62 | 51.82 | 7 | NA* |
| DP4 | ~300y | R-g | SE | 8,579,357 | 0.36 | 0.33 | 0.01 | NA* |
| DP5 | ~300y | R-g | SE | 7,950,283 | 2.55 | 0.93 | 0.07 | NA* |
| DP6 | ~10y | TP | PE | 85,442,043 | 90.53 | 69.21 | 51 | 1,732,886 |
| DP7 | ~10y | TP | PE | 134,031,117 | 90.95 | 69.58 | 78 | 1,734,987 |
| DP8 | ~180y | TP | PE | 95,665,657 | 91.65 | 69.50 | 58 | 1,742,463 |
| DP9 | ~180y | TP | PE | 93,262,683 | 91.85 | 69.55 | 57 | 1,741,613 |

\*no variants called in these sample due to insufficient sequencing depth for triploid variants

**b)**
| Mitochondrial genome *D. pulicaria* |  |  |  |  |  |  |
| --- | --- | --- | --- | --- | --- | --- |
| sample | age | Seq | mapped reads % | coverage breadth % | coverage depth | SNP loci |
| DP1 | ~10y | SE | 73.87 | 99.97 | 61,542 | 200 |
| DP2 | ~140y | SE | 0.68 | 99.94 | 498 | 200 |
| DP3 | ~140y | SE | 0.25 | 99.93 | 252 | 200 |
| DP4 | ~300y | SE | 0.13 | 99.93 | 71 | 200 |
| DP5 | ~300y | SE | 4.55 | 99.95 | 2,315 | 200 |
| DP6 | ~10y | PE | 3.32 | 99.97 | 25,740 | 200 |
| DP7 | ~10y | PE | 0.86 | 99.95 | 10,240 | 200 |
| DP8 | ~180y | PE | 0.72 | 99.97 | 6,155 | 200 |
| DP9 | ~180y | PE | 0.62 | 99.95 | 5,197 | 200 |

*no variants called in these sample due to insufficient sequencing depth for triploid variants

High-throughput sequencing of libraries prepared from WGA-DNA from 13 of the 16 tested dormant *Daphnia* eggs successfully mapped with between 48% and 99% of reads (mean 88%, median 92%) in both SE and PE libraries to the respective *Daphnia* nuclear genomes, suggesting that WGA largely resulted in amplification of the target DNA (Table 2a). This was achieved for ephippia produced in lab cultures as well as those collected from lake sediment of different age with up to ~180y old eggs. Maximum coverage breadth, i.e. the fraction of the nuclear genome covered was similar in both *Daphnia* species (between 70 and 80%, Table 2a). Moderate mapping efficiency was recorded for two relatively young eggs at 48% (DP1, from ~10y old sediment) and 56% (DM4, from a lab culture) and resulted in coverage of a lower fraction of the nuclear reference genomes (lower coverage breadth). Mapping to the nuclear genome failed almost entirely in the two oldest eggs where WGA was attempted (DP4 and DP5, ~300 y old, Table 2a), although the WGA-DNA yield was similar to that of other eggs (Table 1), suggesting amplification of contaminant DNA. However, egg age did not have a consistent effect on mapping efficiency, e.g. libraries for three ~10y old *D. pulicaria* eggs mapped with an efficiency between 48% (DP1) and > 90% (DP6, DP7), while the libraries from two ~140y old eggs were mapped with an efficiency of 84-87% (DP2, DP3).

In contrast to the nuclear genome, we found that reads obtained from all historical eggs of *D. pulicaria* including the oldest samples (~300y old) could be mapped to the mitochondrial genomes of the target species, resulting in high average coverage depth between 71X and 60,000X and a near to 100% coverage breadth of the mitochondrial genome (Table 2b).

### Read distribution

Patterns of genome-wide normalized read distribution differed between species, WGA kits and sequencing strategy (Fig. 1). In *D. magna*, reads originating from the REPLI-g kit (SE sequencing, Fig. 1 A-B) produced an even coverage that was repeatable between samples. Read distribution of *D. magna* DNA obtained from this kit appeared to be more uniform, and to produce fewer and less apparent outliers compared with the TruePrime kit in this species. The read distribution pattern differed markedly from that observed in *D. pulicaria* (Fig. 1 C-E). Samples from this species were represented by historical, sedimentary eggs and therefore the DNA may have been compromised to different degrees, providing inferior amplification substrate. Read distribution obtained from sequencing SE libraries of sedimentary eggs (only *D. pulicaria*, Fig. 1C) appeared less uniform in the REPLI-g amplified samples (DP1 – DP3), which revealed several regions of the genome that were preferentially amplified. These outliers included genomic regions that code e.g. for the Pokey transposon or several introns, but also for segments of mitochondrial DNA encoded in the nuclear genome (Table S1). SE libraries obtained from historical eggs amplified with TruePrime provided a more uniform coverage with less pronounced outliers even in samples > 100y old (Fig. 1C,E). The same observation was made for the PE libraries obtained from TruePrime amplified samples (Fig. 1D,E). The most extreme outlier identified in TruePrime-amplified DNA (a segment of nuclear mitochondrial DNA on scaffold 38, Table S1) was observed in both SE and PE libraries of the historical eggs.

**Fig. 1.**
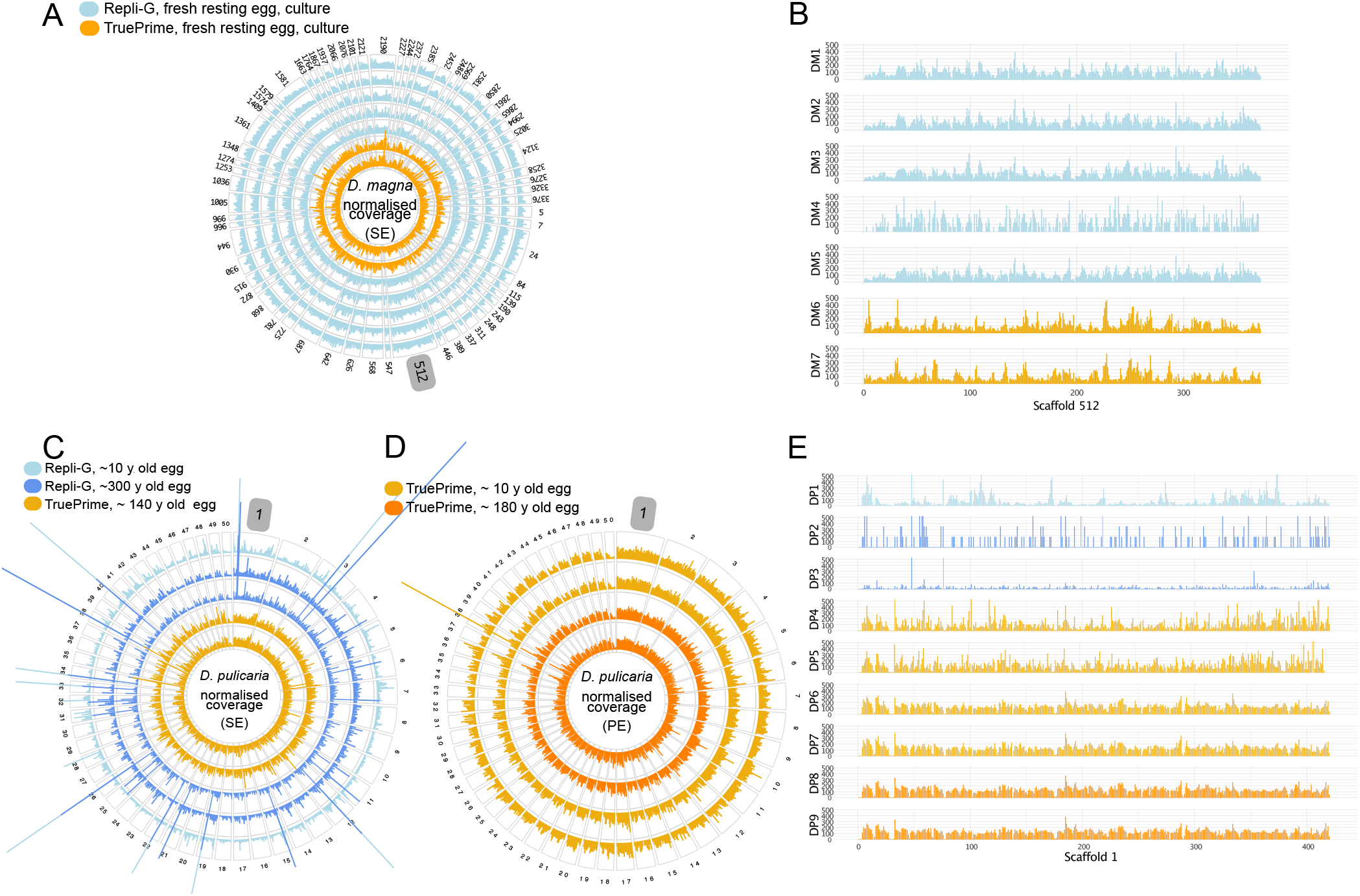
A-E. Visualisation of the genome-wide normalised coverage pattern resulting from WGA-DNA of dormant eggs of *D. magna* and *D. pulicaria*, with comparison of two commercial MDA kits. A) Seven *D. magna* samples (DM1 – DM7, dormant eggs from cultures), amplified with REPLI-g (five outer rings, light blue) or TruePrime (two inner rings, orange). Bars represent normalised coverage (SE sequencing) in 100 kb bins on the N50 scaffolds. B) Detail of A) for the largest *D. magna* scaffold (scaffold 512). Bars represent normalised coverage in 10 kb bins. C) Five *D. pulicaria* samples (DP1 – DP5, eggs isolated from sediment of different age). Amplification with REPLI-g (10 year old egg, light blue, 300 y old egg, blue TruePrime (two inner rings, orange). Bars represent normalised coverage (SE sequencing) in 100 kb bins on the N50 scaffolds. D) Detail of C) for the largest *D. pulex* scaffold (scaffold 1). Bars represent normalised coverage in 10 kb bins

To test repeatability of read distribution patterns between samples within each species, we computed pairwise correlation coefficients (excluding the outlier on scaffold 38) for read counts within 10 kb bins (Fig. 2, Figs. S2, S3). For both species, samples amplified by the same WGA kit were strongly correlated (mean r: *D. magna* REPLI-g = 0.731; *D. magna* TruePrime = 0.767; *D. pulicaria* REPLI-g (SE) = 0.643; *D. pulicaria* TruePrime (SE) = 0.470, *D. pulicaria* TruePrime (PE) = 0.938). Weak correlations were observed between samples amplified using different kits (mean r: *D. magna =* 0.149; *D. pulicaria =* 0.085).

**Fig. 2.**
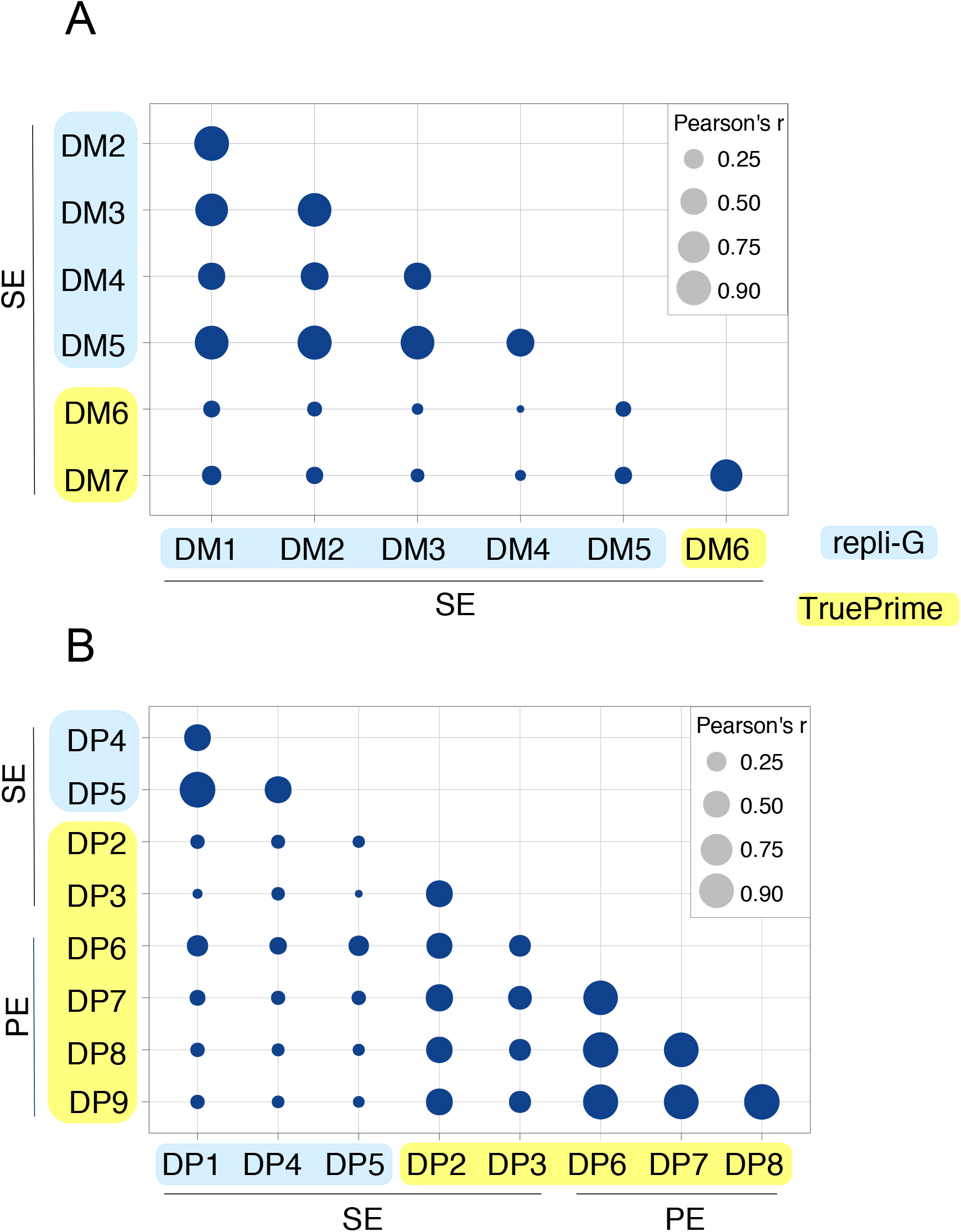
Correlation matrix (Pearson’s r) of pairwise comparisons between normalised read counts of samples in 10 kb bins. A) Pairwise comparisons of *D. magna* samples. B) Pairwise comparisons of *D. pulicaria* samples. All pairwise comparisons had an associated p-value < 0.01

The read distributions of none of the amplified samples had a significant fit to a Poisson distribution expected under the standard model for random sequencing (Table S2).

### Exogenous DNA

In samples where WGA failed almost entirely to produce target DNA (DP4 and DP5, Table 2a), but yielded a similar amount of DNA as other eggs, we used a BLAST search to identify taxon origin of the unmapped reads. This DNA was identified mostly as contaminant DNA from various bacterial and invertebrate taxa, as well as human DNA, but diversity and percentages of the contaminant taxa differed between the two eggs (Fig. 3 and Fig. S4).

**Fig. 3.**
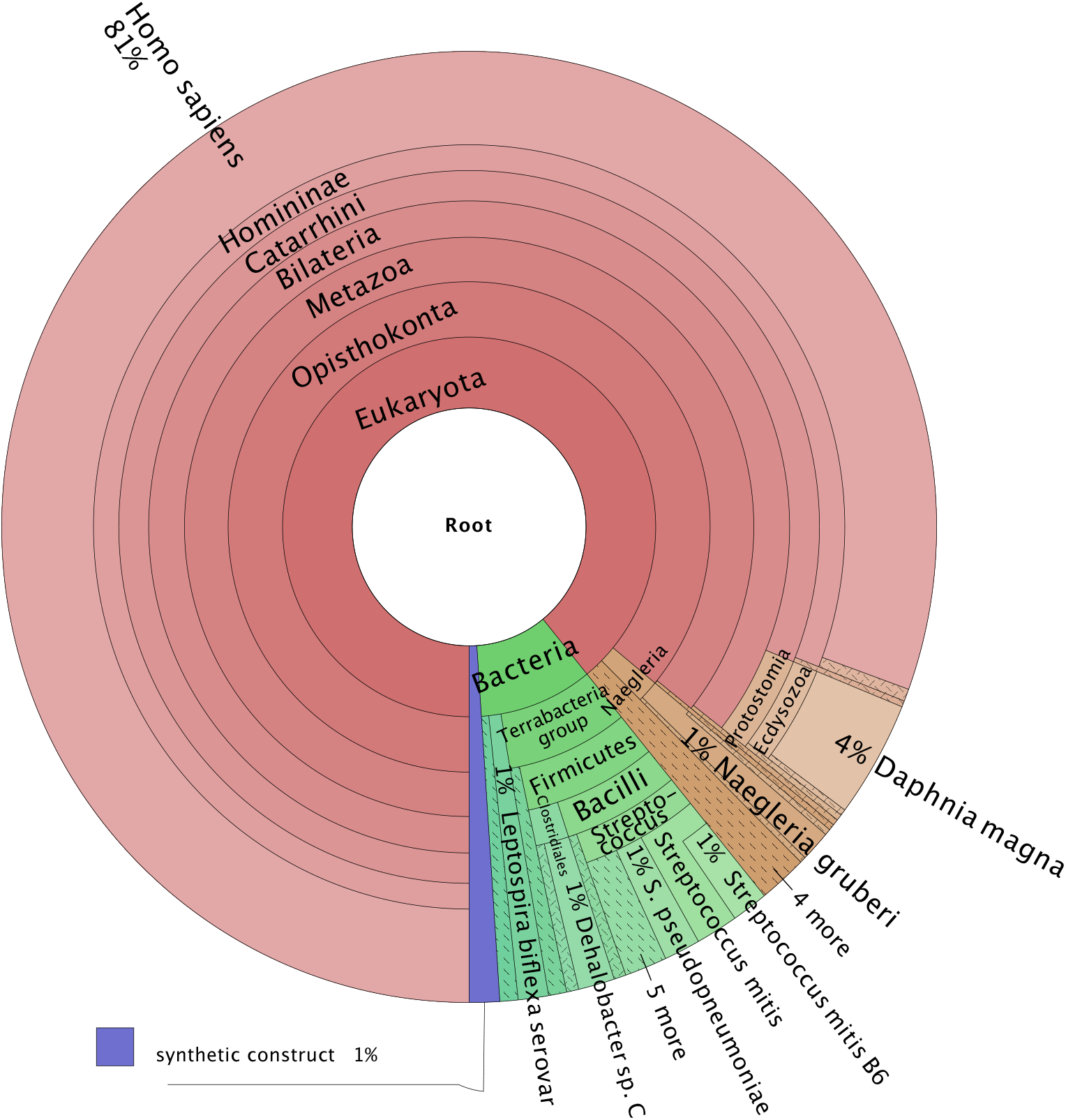
Taxon identity of non-target DNA amplified from a *D. pulicaria* dormant egg (DP5)

### Variant calling

For *D. magna* we called single nucleotide variants (SNPs) in the six samples with > 6 million reads (DM1-DM3, DM5-DM7). For the triploid *D. pulicaria* eggs we could only use the four PE samples for confident SNP calling because deeper sequencing with higher coverage depth is needed for higher ploidy genomes.

The number of identified SNPs per sample in *D. magna* was between 99,927 and 430,909 in comparison with the *D. magna* reference genome (Table 2). In the *D. magna* samples with the highest number of called SNPs (REPLI-g: DM2, DM3, TruePrime: DM7, Table 2, Fig. 4A), a total of 442,976 unique and shared SNP loci were recorded. All three samples could be genotyped at the required depth at 327,802 of these positions (74% of the potential genomic positions of SNPs in these three samples). SNP loci recorded in the lower-depth *D. magna* sample DM6 were nested almost entirely in those of the higher-depth sample DM7 (both amplified with TruePrime, Fig. 4B). In *D. pulicaria*, the total number of SNP loci identified in the four PE samples in comparison to the *D. pulex* reference genome was 1,758,439. All four samples could be genotyped with the required depth at 1,712,889 (or 97.4%) of these genomic positions (Fig. 4C).

**Fig. 4.**
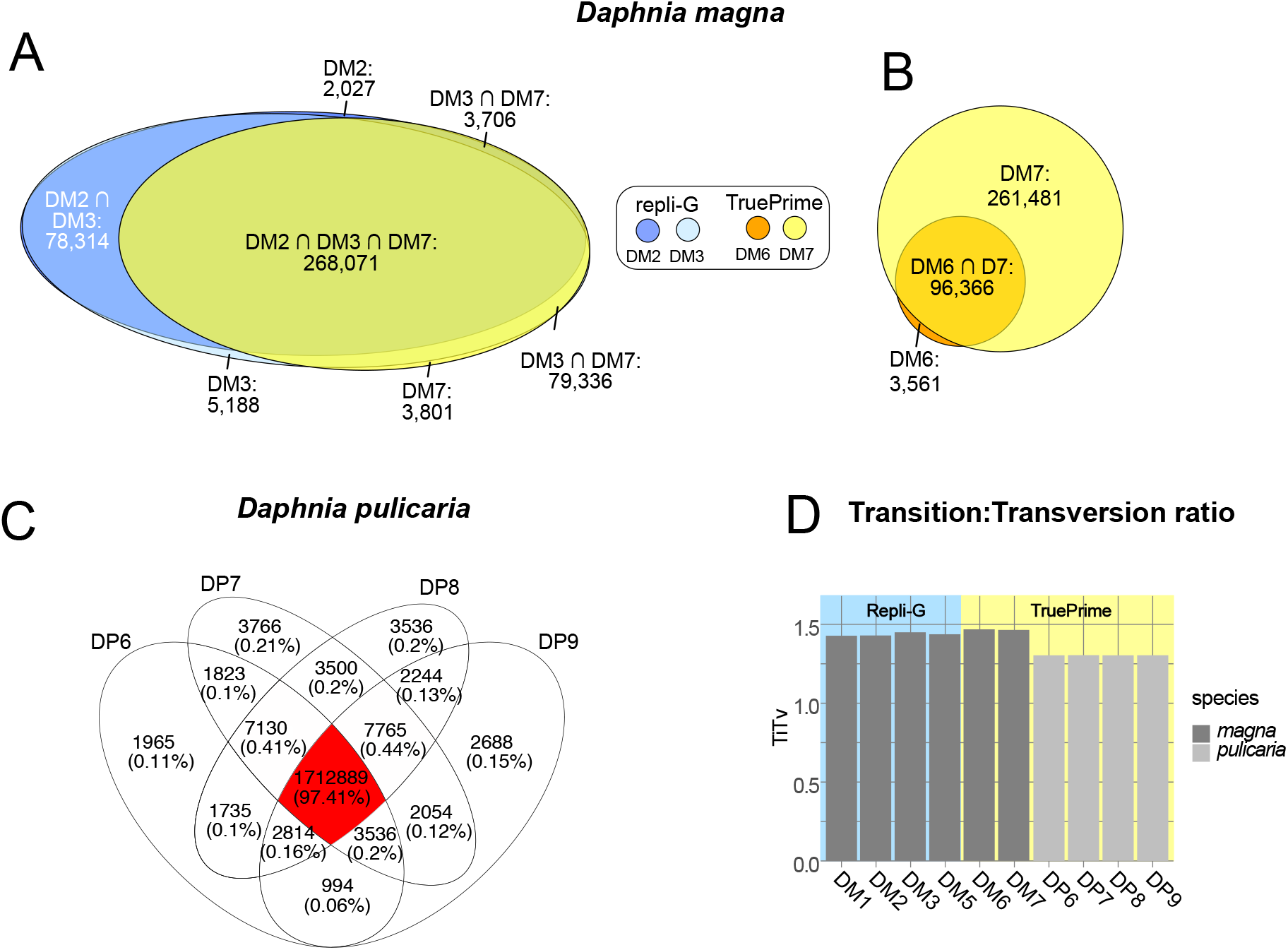
Results of SNP analysis in *D. magna* and *D. pulicaria*. Venn diagrams (A-C) represent the number of genomic positions where SNPs were located in one or more samples and that were sequenced in all or a subset of samples (shared and unique genomic positions). This comparison was used to estimate the repeatability of whole genome amplification and thus for the resulting capacity to call variants at multi-sample level. A) Comparison between three *D. magna* samples with the highest number of SNPs identified, amplified by REPLI-g (DM2, DM3) and TruePrime (DM7). B) Comparison between the two *D. magna* samples amplified by TruePrime (DM6, DM7). C) Comparison between four *D. pulicaria* samples, amplified with TruePrime. D) Transition:transversion ratio analysed for SNPs of both *Daphnia* species

We estimated the transition:transversion ratio (ti:tv, Fig. 4D) to gage the variability of this metric between the samples studied and to assess whether their values were within the ranges reported for *Daphnia* in general. We found that the ratios differed between species but that variation among samples of the same species was small (*D. pulicaria*: 1.30 in all four samples, *D. magna* 1.43 – 1.47). Differences of ti:tv ratios between amplification kits (only *D. magna*) were small (REPLI-g: 1.43 – 1.45, TruePrime: 1.46 – 1.47).

SNP calling in the mitochondrial genomes (Fig. 5) was performed with all available *D. pulicaria* samples. The analysis revealed the presence of 200 biallelic SNPs that differed between the two lake populations sampled. However, within-populations samples were identical with the exception of DP1 (Lake SS4) and DP9 (Lake 1381) that differed from other samples of their respective population by a single SNP.

**Fig. 5.**
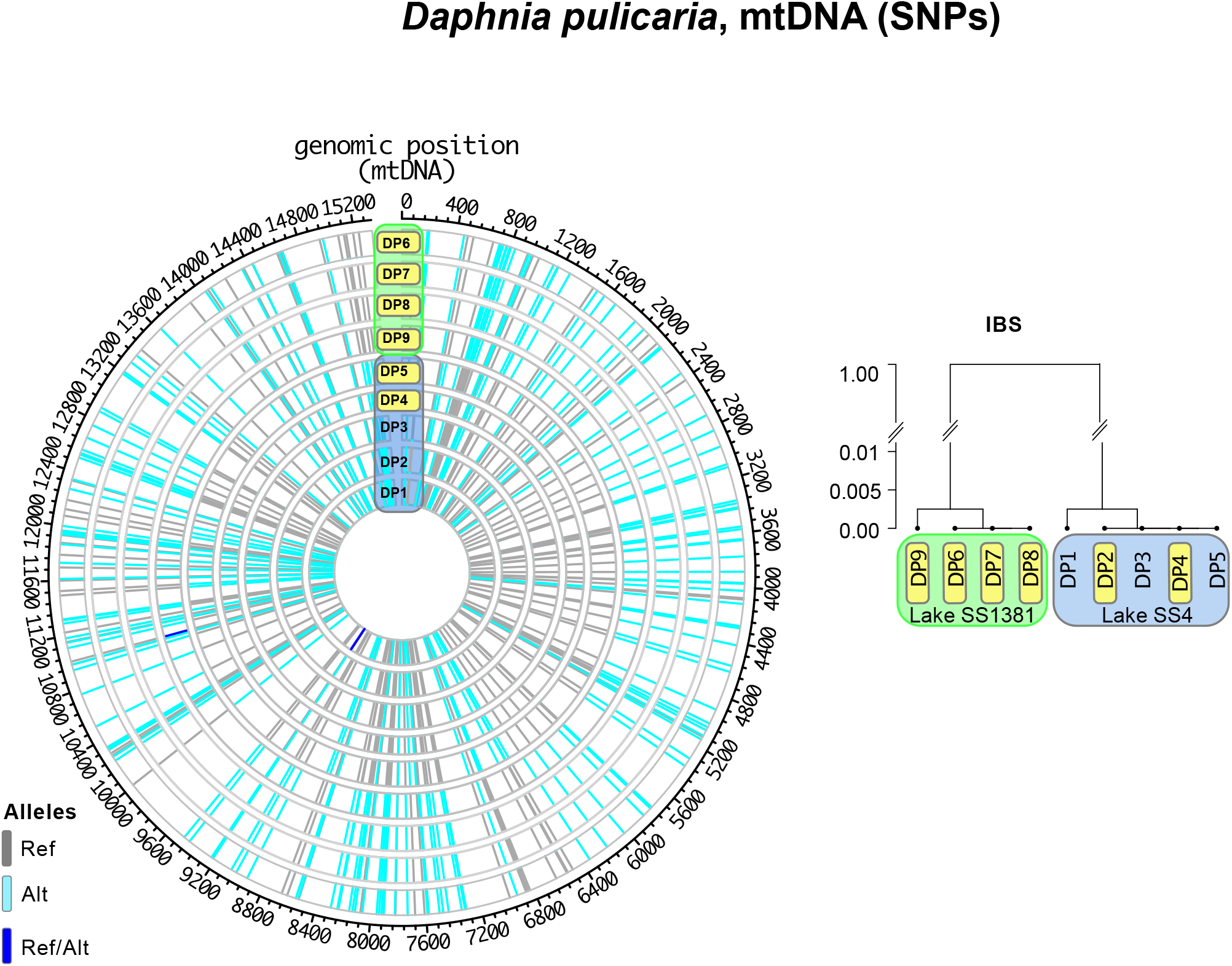
SNP positions and pairwise clustering of individuals by Identity-by-state (IBS) of nine *D. pulicara* mitochondrial genomes recovered from historical sedimentary dormant eggs from two lakes in West Greenland (Lake SS1381, green label, and SS4, blue label). Samples with yellow labels were amplified with TruePrime, and the remaining three with REPLI-g. A) Circular plot of detected SNP loci. The four outer rings represent samples from Lake SS1381, the five inner rings from SS4. SNP colours are labelled according to their state (turquoise = homozygous (variant allele), grey = homozygous (reference allele), blue = heterozygous (both reference and variant allele present). B) Hierarchical clustering tree resulting from the fraction of identical genomic positions (Identity-by-State) in the nine mitochondrial genomes compared

## Discussion

Whole genome amplification and subsequent whole genome sequencing are staples of modern single-cell genome studies, and are widely applied to the study of human diseases (Huang, Ma, Chapman, Lu, & Xie, 2015). Other promising but less common applications include phylogenomics (Ahrendt et al., 2018; Zhang et al., 2019) and metagenomics of microbial communities (Xu & Zhao, 2018). Suitable application for population genomics and evolutionary studies using *Daphnia* dormant eggs has been suggested (Lack et al., 2018), but to date a comprehensive study involving the comparison between multiple sedimentary eggs of different historical age and species, applying different pre-treatments and amplification kits is not available. Multiple displacement amplification (MDA) has superior qualities when the goal is the discovery of single nucleotide variants, due to high fidelity of the φ29 polymerase and associated low error rates, while PCR-based WGA such as MALBAC may perform better for detecting copy number variation (Chen et al., 2014; de Bourcy et al., 2014). We therefore tested two MDA-WGA kits to provide a detailed workflow (Fig. 6) for successful and repeatable amplification and sequencing of DNA for variant calls from dormant eggs of *Daphnia*.

**Fig. 6.**
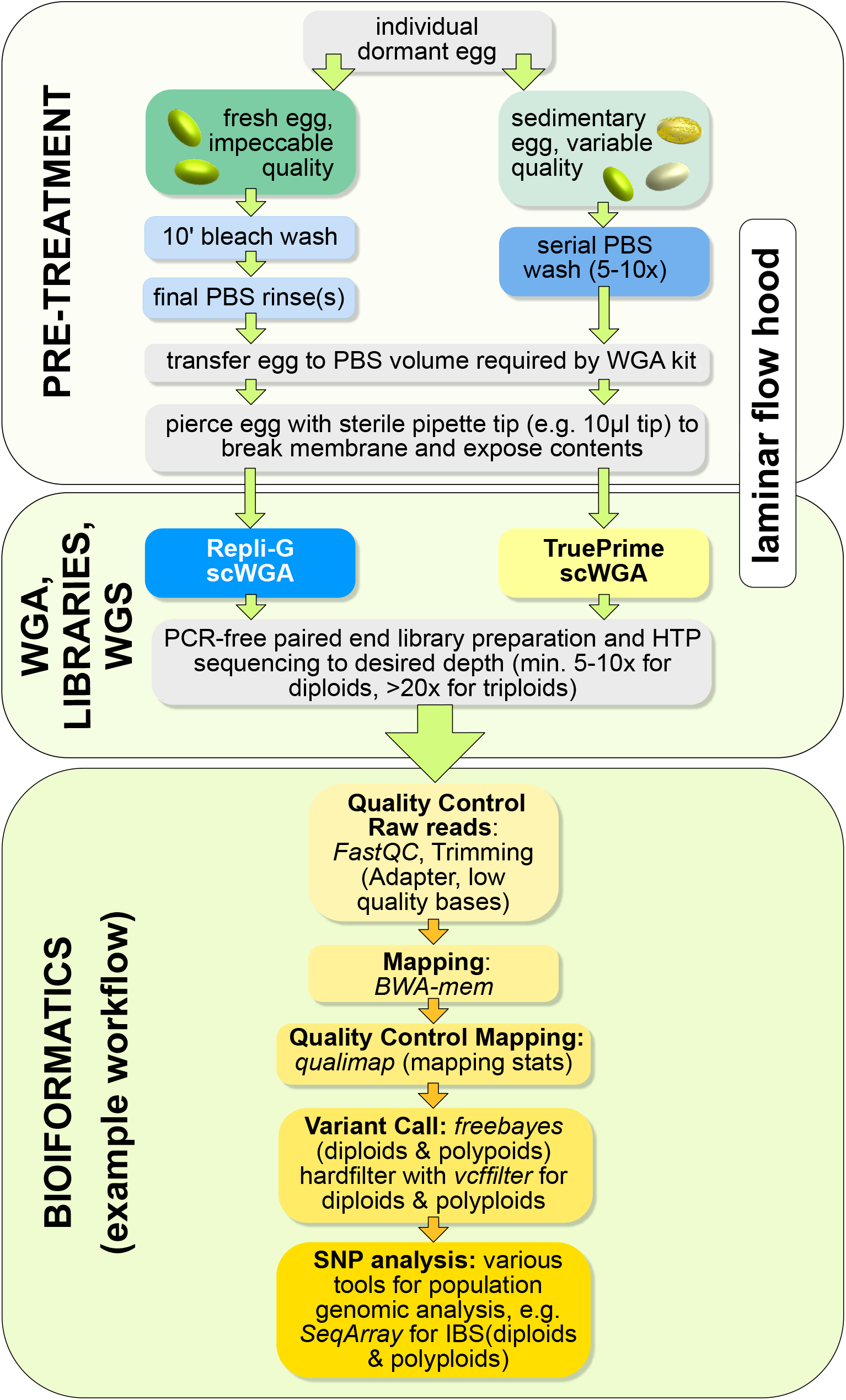
Recommended workflow for whole genome amplification and whole genome sequencing with downstream variant call and variant filtering. Recommendation for sequencing depth of higher ploidy genomes see (Maruki & Lynch, 2017). For further details see text

### Pre-Treatment to minimise exogenous DNA

Sources of DNA contamination may originate from the WGA reagents, or are introduced when handling samples (Rinke et al., 2014). Contamination may also derive from non-target exogenous DNA on the biological isolate (often of microbial origin), which is especially common in historical samples containing highly degraded ancient DNA (Pilli et al., 2013). Decontamination procedures of equipment, which include the application of bleach and UV light, can be performed prior to WGA to prevent the amplification of exogenous DNA (Weiss *et al.*, 2014). It is also recommended to use a thoroughly decontaminated laminar flowhood during all stages of WGA, preferably situated in a dedicated clean room.

Overall, our data suggests that amplification of exogenous DNA can be kept to a minimum when all careful steps are followed to avoid contamination. However, in historical samples where DNA is already damaged, exogenous DNA may be present in higher amounts than the target DNA, and thus be preferentially amplified, and even overwhelm the amplification process. This was particularly obvious in the oldest samples tested here (~300y old). Apart from egg age, DNA integrity may also be highly dependent on the preservation conditions of the lake sediment: DNA preservation can vary strongly between different lakes and may be related to a number of variables, including temperature, salt concentration and pH conditions (Ellegaard et al., 2020; Giguet-Covex et al., 2019)

Washes with diluted bleach of *D. pulicaria* sedimentary eggs had detrimental impact on DNA integrity of both younger and older eggs (~10 to 300y old) and therefore on the quality of WGA-DNA. In contrast, washing with PBS did not appear to affect the DNA in dormant eggs of *D. pulicaria* of different historical age (~10 and ~180y old), which produced high quality WGA-DNA. It is also possible that these eggs had a generally higher quality than those used for bleaching, which were sampled from sediment of a neighbouring lake. Nevertheless, all sedimentary *D. pulicaria* eggs were older than the dormant eggs removed from laboratory cultures of *D. magna* in our study. These were only several days old and did not show any signs of DNA degradation even after bleaching with a higher bleach concentration, and/or for a longer exposure time to bleach. A possible explanation is the likely presence of microfissures in the egg membranes of historical, sedimentary eggs that are prone to increase with dormant egg age. To test this idea, microscopic studies comparing eggs of different age are needed. Based on these findings, we recommend the use of careful serial PBS washes instead of diluted bleach to remove possible contaminants on sedimentary egg surfaces that could overwhelm DNA amplification, particularly if egg DNA is more strongly impaired with increasing age.

Mapping of WGA-DNA produced with either of the two kits tested here was highly successful in most isolates, with a median of 92% of reads mapped to their respective reference genomes, even of eggs as old as 180 years. Coverage breadth in most samples was between 70-80%. These values are similar to that previously reported for a dormant *Daphnia* egg, but below those obtained from *Daphnia* bulk sequencing (Lack et al., 2018), and higher than of MALBAC amplified individual sperm cells of *Daphnia* (53%, Xu et al., 2015). Indeed, incomplete genome coverage is commonly observed in WGA data in various taxa (e.g. de Bourcy et al., 2014; Huang et al., 2015; Picher et al., 2016).

Deviation of read distributions from a Poisson distribution expected under a random sequencing model (Lander & Waterman, 1988) was observed in all samples. However, this is not unexpected as it has been described previously that this model is inadequate for single-cell sequencing due to the possibility of locus dropout (Daley & Smith, 2014). Despite this, we found the regions of the genome that are amplified to be remarkably repeatable among samples of the same species, with highly significant correlation coefficients within amplification kits (average r of ~0.7). The r values below average were generally associated with failed target amplification or low read depth overall, specifically in the SE samples, but above average in the PE samples with high read depth. However, correlations between WGA kits were not strong, so for good comparability between samples, it is recommended to apply only one kit.

The highly reproducible patterns of read distribution within kits are largely responsible for the success of SNP calling across samples, demonstrating the suitability of this method for population genomic applications particularly for dormant eggs and thus for its utility for studying genome evolution using sediment archives. In *D. magna*, >70% of the SNP positions could be called in all three samples with >14 million reads. Perhaps not surprisingly these values were even higher in the PE samples of *D. pulicaria* with >80 million reads, suggesting in general that sequencing strategy and depth strongly influence fidelity of the variant call also when applied to WGA-DNA.

Transition to transversion ratios have been suggested as a quality indicator for human SNP discovery (Wang, Raskin, Samuels, Shyr, & Guo, 2015). However, because these ratios vary between species, e.g. averaging 1.54 in several strains of *Daphnia magna* (Ho et al., 2020), or 0.45 in *C. elegans* (Denver et al., 2009), comparisons should be made within species. Our range between 1.43 to 1.47 for nuclear DNA of *D. magna* was well within the ratio measured by Ho et al. (2020), and for *D. pulicaria* (ti:tv 1.30) was similar to that reported for the closely related *D. pulex* (1.58, Keith et al., 2016). In this study, we can also apply the ti:tv ratio as indicator for the reliability of the WGA procedure within and across kits, with highly similar values for *D. magna* eggs, or almost identical values for *D. pulicaria*.

Other studies have identified possible limitations of the reliability of WGA, such as coverage uniformity, reproducibility, and allele dropout rate (de Bourcy et al., 2014; Huang et al., 2015). The substrates that these studies used were cell lineages from which individual cells were subjected to scWGA and compared to bulk sequencing of a multicellular sample from the same lineage. For historical isolates of dormant eggs from the sediment, such a strategy cannot be applied. However, an effective substitute to test the reliability of SNPs from these historical samples was the use of asexually produced dormant eggs such as those of the Arctic *D. pulicaria* population from which our samples originated. WGA of these samples allowed the genotyping of 97% of all SNP loci detected in the PE sequences. Preliminary analysis of variation between these genotyped eggs showed a maximum difference of 3% of the roughly 1.7 million SNPs between individuals (unpublished data).

An added benefit of WGA with both tested kits was the possibility of obtaining high coverage of the full mitochondrial genome for both species. This is of particular interest for the historical samples from which full mitochondrial genomes and high-quality SNP calls could be retrieved even of the oldest samples (~300y old eggs). This opens a promising avenue for gathering information of genome-wide mutation rates and spectra of the mitochondrial genome stored in sedimentary archives across extended time periods and thousands of generations, likely surpassing the time range tested here.

## Conclusion

Although the method described here was tested on dormant eggs of *Daphnia*, it could be widely applied to sedimentary dormant stages of a variety of taxa, potentially providing access not only to single taxa but also to entire aquatic communities. Such taxa could include key members of the plankton in the freshwater and marine to hypersaline ecosystems that produce dormant stages, such as the dormant eggs of calanoid copepods, cysts of the brine shrimp *Artemia*, algal dormant stages and plant seeds. Other methods that have successfully been applied to individual small-sized planktonic crustaceans, such as low-DNA-input sequencing libraries (Beninde, Möst, & Meyer, 2020) could be tested on dormant stages isolated from historical sediment layers. However their protocol, which was optimised for adult specimens requires a highly efficient DNA extraction step and includes up to 8 PCR cycles to obtain library concentrations as high as 5 ng DNA. Since DNA concentration in eggs or other propagules is generally very low, a DNA extraction step which inevitably leads to a certain amount of DNA loss, could be imprudent when precious historical material is involved.

Our data reveal that both of the tested amplification kits provided high quality DNA for most *Daphnia* egg isolates, and that the amplified DNA could efficiently be applied to WGS and subsequent genome-wide studies at the population level. However, differences were observed with respect to the age of dormant eggs, where our results suggest a superior performance of TruePrime (compared with REPLI-g) for application to eggs of sedimentary origin and thus to prospectively degraded DNA. A possible explanation could be that the primase TthPrimPol used in the TruePrime kit shows translesion activity, allowing reinitiation of the replication fork when encountering damaged DNA, and thus continued amplification of damaged substrate (Picher & Blanco, 2014). Due to the similarity of TthPrimPol to human PrimPol, another mechanism could explain our results: human PrimPol can reprime DNA synthesis following a lesion, allowing the φ29 DNA polymerase to continue the amplification process close to the region in which the lesion was found (Mourón et al., 2013), possibly improving evenness during the process.

Ultimately, our data indicate that for optimal results, preliminary trials are recommended using both kits tested here (and if possible others) on the dormant egg population in question.

## Acknowledgements

DF received funding from the European Union’s Horizon 2020 research and innovation programme under the Marie Skłodowska-Curie grant agreement No. 658714 and NERC Biomolecular Analysis Facility Pilot Project Grant NBAF998. COG received funding by the Midlands Integrative Biosciences Training Partnership (MIBTP). We are grateful to Stephen Kissane for preparation and sequencing of single-end libraries, and to Caroline Sewell for supplying *D. magna* ephippia from the *Daphnia* facility, UoB. Paired-end library preparation and sequencing were carried out by Edinburgh Genomics, the University of Edinburgh, which is partly supported with core funding from NERC (UKSBS PR18037).

## Data accessibility and Benefit-Sharing Statement

The sequence data supporting the findings reported here will be made available in a suitable public data repository such as NCBI Genbank or Dryad upon acceptance of the manuscript.

## Author contributions

DF conceived the idea and obtained funding for the study. DF, COG and JKC designed the experiments. COG performed the labwork. DF and COG analysed the data and wrote the paper with input from JKC and VD.

## References

Ahrendt, S. R., Quandt, C. A., Ciobanu, D., Clum, A., Salamov, A., Andreopoulos, B.,…Grigoriev, I. V. (2018). Leveraging single-cell genomics to expand the fungal tree of life. Nature Microbiology, 3(12), 1417–1428. doi: 10.1038/s41564-018-0261-0

Andrews, S. (2015). FASTQC A Quality Control tool for High Throughput Sequence Data. Babraham Institute. Https://Www.Bioinformatics.Babraham.Ac.Uk/Projects/Fastqc/.

Beninde, J., Möst, M., & Meyer, A. (2020). Optimized and affordable high-throughput sequencing workflow for preserved and nonpreserved small zooplankton specimens. Molecular Ecology Resources, 20(6), 1632–1646. doi: 10.1111/1755-0998.13228

Blainey, P. C., & Quake, S. R. (2011). Digital MDA for enumeration of total nucleic acid contamination. Nucleic Acids Research. doi: 10.1093/nar/gkq1074

Blanco, L., Bernad, A., Lázaro, J. M., Martín, G., Garmendia, C., & Salas, M. (1989). Highly Efficient DNA Synthesis by the Phage ϕ 29 DNA Polymerase. Journal of Biological Chemistry. doi: 10.1016/s0021-9258(18)81883-x

Bolger, A. M., Lohse, M., & Usadel, B. (2014). Trimmomatic: A flexible trimmer for Illumina sequence data. Bioinformatics. doi: 10.1093/bioinformatics/btu170

Brede, N., Sandrock, C., Straile, D., Spaak, P., Jankowski, T., Streit, B., & Schwenk, K. (2009). The impact of human-made ecological changes on the genetic architecture of *Daphnia* species. Proceedings of the National Academy of Sciences of the United States of America, 106(12), 4758–4763.

Broad Institute. (2019). Picard toolkit. Broad Institute, GitHub Repository. Https://Github.Com/Broadinstitute/Picard/Releases/Tag/2.25.0.

Chen, L., Barnett, R. E., Horstmann, M., Bamberger, V., Heberle, L., Krebs, N.,…Weiss, L. (2018). Mitotic activity patterns and cytoskeletal changes throughout the progression of diapause developmental program in *Daphnia*. BMC Cell Biology, 19(1), 30. doi: 10.1186/s12860-018-0181-0

Chen, M., Song, P., Zou, D., Hu, X., Zhao, S., Gao, S., & Ling, F. (2014). Comparison of Multiple Displacement Amplification (MDA) and Multiple Annealing and Looping-Based Amplification Cycles (MALBAC) in Single-Cell Sequencing. PLoS ONE, 9(12), e114520. doi: 10.1371/journal.pone.0114520

Cheung, V. G., & Nelson, S. F. (1996). Whole genome amplification using a degenerate oligonucleotide primer allows hundreds of genotypes to be performed on less than one nanogram of genomic DNA. Proceedings of the National Academy of Sciences, 93(25), 14676–14679. doi: 10.1073/pnas.93.25.14676

Colbourne, J. K., Pfrender, M. E., Gilbert, D., Thomas, W. K., Tucker, A., Oakley, T. H.,…Boore, J. L. (2011). The Ecoresponsive Genome of *Daphnia pulex*. Science, 331(6017), 555–561. doi: 10.1126/science.1197761

Cordellier, M., Wojewodzic, M. W., Wessels, M., Kuster, C., & von Elert, E. (2021). Next-generation sequencing of DNA from resting eggs: signatures of eutrophication in a lake’s sediment. Zoology. doi: 10.1016/j.zool.2021.125895

Crease, T. J. (1999). The complete sequence of the mitochondrial genome of *Daphnia pulex* (Cladocera: Crustacea). Gene. doi: 10.1016/S0378-1119(99)00151-1

Daley, T., & Smith, A. D. (2014). Modeling genome coverage in single-cell sequencing. Bioinformatics. doi: 10.1093/bioinformatics/btu540

Dane, M., Anderson, N. J., Osburn, C. L., Colbourne, J. K., & Frisch, D. (2020). Centennial clonal stability of asexual *Daphnia* in Greenland lakes despite climate variability. Ecology and Evolution, 10(24), 14178–14188. doi: 10.1002/ece3.7012

de Bourcy, C. F. A., De Vlaminck, I., Kanbar, J. N., Wang, J., Gawad, C., & Quake, S. R. (2014). A quantitative comparison of single-cell whole genome amplification methods. PloS One, 9(8), e105585. doi: 10.1371/journal.pone.0105585

De La Torre, A. R., Wilhite, B., & Neale, D. B. (2019). Environmental Genome-Wide Association Reveals Climate Adaptation Is Shaped by Subtle to Moderate Allele Frequency Shifts in Loblolly Pine. Genome Biology and Evolution, 11(10), 2976–2989. doi: 10.1093/gbe/evz220

Dean, F. B., Hosono, S., Fang, L., Wu, X., Faruqi, A. F., Bray-Ward, P.,…Lasken, R. S. (2002). Comprehensive human genome amplification using multiple displacement amplification. Proceedings of the National Academy of Sciences of the United States of America, 99(8), 5261–5266. doi: 10.1073/pnas.082089499

Dean, F. B., Nelson, J. R., Giesler, T. L., & Lasken, R. S. (2001). Rapid amplification of plasmid and phage DNA using Phi29 DNA polymerase and multiply-primed rolling circle amplification. Genome Research. doi: 10.1101/gr.180501

Denver, D. R., Dolan, P. C., Wilhelm, L. J., Sung, W., Lucas-Lledo, J. I., Howe, D. K.,…Baer, C. F. (2009). A genome-wide view of *Caenorhabditis elegans* base-substitution mutation processes. Proceedings of the National Academy of Sciences, 106(38), 16310–16314. doi: 10.1073/pnas.0904895106

Dettman, J. R., Rodrigue, N., Melnyk, A. H., Wong, A., Bailey, S. F., & Kassen, R. (2012). Evolutionary insight from whole-genome sequencing of experimentally evolved microbes. Molecular Ecology. doi: 10.1111/j.1365-294X.2012.05484.x

Dorant, Y., Benestan, L., Rougemont, Q., Normandeau, E., Boyle, B., Rochette, R., & Bernatchez, L. (2019). Comparing Pool-seq, Rapture, and GBS genotyping for inferring weak population structure: The American lobster (*Homarus americanus*) as a case study. Ecology and Evolution, 9(11), 6606–6623. doi: 10.1002/ece3.5240

Ellegaard, M., Clokie, M. R. J., Czypionka, T., Frisch, D., Godhe, A., Kremp, A.,…John Anderson, N. (2020). Dead or alive: sediment DNA archives as tools for tracking aquatic evolution and adaptation. Communications Biology, 3(1), 169. doi: 10.1038/s42003-020-0899-z

Ellegren, H. (2014). Genome sequencing and population genomics in non-model organisms. Trends in Ecology and Evolution. doi: 10.1016/j.tree.2013.09.008

Frisch, D., Morton, P. K., Culver, B. W., Edlund, M. B., Jeyasingh, P. D., & Weider, L. J. (2016). Paleogenetic records of *Daphnia pulicaria* in North American lakes reveal the impact of cultural eutrophication. Global Change Biology, 23(2), 708–718. doi: 10.1111/gcb.13445

Frisch, D., Morton, P. K., Roy Chowdhury, P., Culver, B. W., Colbourne, J. K., Weider, L. J., & Jeyasingh, P. D. (2014). A millennial-scale chronicle of evolutionary responses to cultural eutrophication in *Daphnia*. Ecology Letters, 17(3), 360–368.

Gao, C.-H., & Yi, L. (2019). ggVennDiagram: A “ggplot2” implement of Venn Diagram. https://github.com/gaospecial/ggVennDiagram.

García-Alcalde, F., Okonechnikov, K., Carbonell, J., Cruz, L. M., Götz, S., Tarazona, S.,…Conesa, A. (2012). Qualimap: evaluating next-generation sequencing alignment data. Bioinformatics, 28(20), 2678–2679. doi: 10.1093/bioinformatics/bts503

Garrison, E. (2016). Vcflib, a simple C++ library for parsing and manipulating VCF files. Retrieved from https://github.com/vcflib/vcflib.

Garrison, E., & Marth, G. (2012). Haplotype-based variant detection from short-read sequencing. ArXiv Preprint ArXiv:1207.3907.

Giguet-Covex, C., Ficetola, G. F., Walsh, K., Poulenard, J., Bajard, M., Fouinat, L.,…Arnaud, F. (2019). New insights on lake sediment DNA from the catchment: importance of taphonomic and analytical issues on the record quality. Scientific Reports, 9(1), 14676. doi: 10.1038/s41598-019-50339-1

Gu, Z. (2014). circlize implements and enhances circular visualization in R. Bioinformatics, 30(19), 2811–2812. doi: 10.1093/bioinformatics/btu393

Handyside, A. H., Robinson, M. D., Simpson, R. J., Omar, M. B., Shaw, M. A., Grudzinskas, J. G., & Rutherford, A. (2004). Isothermal whole genome amplification from single and small numbers of cells: A new era for preimplantation genetic diagnosis of inherited disease. Molecular Human Reproduction. doi: 10.1093/molehr/gah101

Härnström, K., Ellegaard, M., Andersen, T. J., & Godhe, A. (2011). Hundred years of genetic structure in a sediment revived diatom population. Proceedings of the National Academy of Sciences of the United States of America, 108(10), 4252–4257.

Ho, E. K. H., Macrae, F., Latta, L. C., McIlroy, P., Ebert, D., Fields, P. D.,…Schaack, S. (2020). High and Highly Variable Spontaneous Mutation Rates in *Daphnia*. Molecular Biology and Evolution, 37(11), 3258–3266. doi: 10.1093/molbev/msaa142

Hohenlohe, P. A., Hand, B. K., Andrews, K. R., & Luikart, G. (2018). Population Genomics Provides Key Insights in Ecology and Evolution. doi: 10.1007/13836_2018_20

Huang, L., Ma, F., Chapman, A., Lu, S., & Xie, X. S. (2015). Single-Cell Whole-Genome Amplification and Sequencing: Methodology and Applications. Annual Review of Genomics and Human Genetics, 16(1), 79–102. doi: 10.1146/annurev-genom-090413-025352

Keith, N., Tucker, A. E., Jackson, C. E., Sung, W., Lucas Lledó, J. I., Schrider, D. R.,…Lynch, M. (2016). High mutational rates of large-scale duplication and deletion in *Daphnia pulex*. Genome Research, 26(1), 60–69. doi: 10.1101/gr.191338.115

Lack, J. B., Weider, L. J., & Jeyasingh, P. D. (2018). Whole genome amplification and sequencing of a *Daphnia* resting egg. Molecular Ecology Resources. doi: 10.1111/1755-0998.12720

Lander, E. S., & Waterman, M. S. (1988). Genomic mapping by fingerprinting random clones: A mathematical analysis. Genomics, 2(3), 231–239. doi: 10.1016/0888-7543(88)90007-9

Larsson, J. (2020). eulerr: Area-Proportional Euler and Venn Diagrams with Ellipses. R package version 6.1.0. Retrieved from https://cran.r-project.org/package=eulerr

Lasken, R. S., & Egholm, M. (2003). Whole genome amplification: abundant supplies of DNA from precious samples or clinical specimens. Trends in Biotechnology, 21(12), 531–535. doi: 10.1016/j.tibtech.2003.09.010

Leonardi, M., Librado, P., Der Sarkissian, C., Schubert, M., Alfarhan, A. H., Alquraishi, S. A.,…Orlando, L. (2017). Evolutionary Patterns and Processes: Lessons from Ancient DNA. Systematic Biology, 66(1), e1–e29. doi: 10.1093/sysbio/syw059

Li, H., & Durbin, R. (2009). Fast and accurate short read alignment with Burrows-Wheeler transform. Bioinformatics, 25(14), 1754–1760. doi: 10.1093/bioinformatics/btp324

Limburg, P. A., & Weider, L. J. (2002). “Ancient” DNA in the resting egg bank of a microcrustacean can serve as a palaeolimnological database. Proceedings of the Royal Society of London Series B-Biological Sciences, 269(1488), 281–287.

Luo, R., Liu, B., Xie, Y., Li, Z., Huang, W., Yuan, J.,…Wang, J. (2012). SOAPdenovo2: An empirically improved memory-efficient short-read de novo assembler. GigaScience. doi: 10.1186/2047-217X-1-18

Maruki, T., & Lynch, M. (2017). Genotype Calling from Population-Genomic Sequencing Data. G3: Genes|Genomes|Genetics, 7(5), 1393–1404. doi: 10.1534/g3.117.039008

Mergeay, J., Verschuren, D., & De Meester, L. (2006). Invasion of an asexual American water flea clone throughout Africa and rapid displacement of a native sibling species. Proceedings of the Royal Society B-Biological Sciences, 273(1603), 2839–2844.

Mourón, S., Rodriguez-Acebes, S., Martínez-Jiménez, M. I., García-Gómez, S., Chocrón, S., Blanco, L., & Méndez, J. (2013). Repriming of DNA synthesis at stalled replication forks by human PrimPol. Nature Structural & Molecular Biology, 20(12), 1383–1389. doi: 10.1038/nsmb.2719

Ondov, B. D., Bergman, N. H., & Phillippy, A. M. (2011). Interactive metagenomic visualization in a Web browser. BMC Bioinformatics, 12(1), 385. doi: 10.1186/1471-2105-2-385

Orsini, L., Schwenk, K., De Meester, L., Colbourne, J. K., Pfrender, M. E., & Weider, L. J. (2013). The evolutionary time machine: forecasting how populations can adapt to changing environments using dormant propagules. Trends in Ecology & Evolution, 28, 274–282.

Orsini, L., Spanier, K. I., & De Meester, L. (2012). Genomic signature of natural and anthropogenic stress in wild populations of the waterflea Daphnia magna: validation in space, time and experimental evolution. Molecular Ecology, 21(9), 2160–2175.

Paez, J. G. (2004). Genome coverage and sequence fidelity of 29 polymerase-based multiple strand displacement whole genome amplification. Nucleic Acids Research, 32(9), e71–e71. doi: 10.1093/nar/gnh069

Parks, M., Subramanian, S., Baroni, C., Salvatore, M. C., Zhang, G., Millar, C. D., & Lambert, D. M. (2015). Ancient population genomics and the study of evolution. Philosophical Transactions of the Royal Society B: Biological Sciences. doi: 10.1098/rstb.2013.0381

Picher, Á. J., & Blanco, L. (2014). Patent No. International Publication Number WO2014140309A1. https://patents.google.com/patent/WO2014140309A1/en

Picher, Á. J., Budeus, B., Wafzig, O., Krüger, C., García-Gómez, S., Martínez-Jiménez, M. I.,…Schneider, A. (2016). TruePrime is a novel method for whole-genome amplification from single cells based on TthPrimPol. Nature Communications, 7(1), 13296. doi: 10.1038/ncomms13296

Pilli, E., Modi, A., Serpico, C., Achilli, A., Lancioni, H., Lippi, B.,…Caramelli, D. (2013). Monitoring DNA Contamination in Handled vs. Directly Excavated Ancient Human Skeletal Remains. PLoS ONE, 8(1), e52524. doi: 10.1371/journal.pone.0052524

Pollard, H. G., Colbourne, J. K., & Keller, W. (2003). Reconstruction of centuries-old *Daphnia* communities in a lake recovering from acidification and metal contamination. Ambio, 32(3), 214–218. doi: 10.1579/0044-7447-32.3.214

Quinlan, A. R., & Hall, I. M. (2010). BEDTools: a flexible suite of utilities for comparing genomic features. Bioinformatics, 26(6), 841–842. doi: 10.1093/bioinformatics/btq033

R Core Team. (2019). R: A Language and Environment for Statistical Computing. R Foundation for Statistical Computing. Vienna, Austria. http://www.R-project.org.

Rajpurohit, S., Gefen, E., Bergland, A. O., Petrov, D. A., Gibbs, A. G., & Schmidt, P. S. (2018). Spatiotemporal dynamics and genome-wide association genome-wide association analysis of desiccation tolerance in *Drosophila melanogaster*. Molecular Ecology. doi: 10.1111/mec.14814

Rinke, C., Lee, J., Nath, N., Goudeau, D., Thompson, B., Poulton, N.,…Woyke, T. (2014). Obtaining genomes from uncultivated environmental microorganisms using FACS-based single-cell genomics. Nature Protocols, 9(5), 1038–1048. doi: 10.1038/nprot.2014.067

Rizzi, E., Lari, M., Gigli, E., De Bellis, G., & Caramelli, D. (2012). Ancient DNA studies: New perspectives on old samples. Genetics Selection Evolution. doi: 10.1186/1297-686-44-21

Sella, G., & Barton, N. H. (2019). Thinking about the Evolution of Complex Traits in the Era of Genome-Wide Association Studies. Annual Review of Genomics and Human Genetics. doi: 10.1146/annurev-genom-083115-022316

Stiller, J., & Zhang, G. (2019). Comparative Phylogenomics, a Stepping Stone for Bird Biodiversity Studies. Diversity. doi: 10.3390/d11070115

Telenius, H., Carter, N. P., Bebb, C. E., Nordenskjöld, M., Ponder, B. A. J., & Tunnacliffe, A. (1992). Degenerate oligonucleotide-primed PCR: General amplification of target DNA by a single degenerate primer. Genomics. doi: 10.1016/0888-7543(92)90147-K

von Baldass, F. (1941). Entwicklung von *Daphnia pulex*. Zoologische Jahrbücher. Abteilung Für Anatomie Und Ontogenie Der Tiere, 67, 1–60.

Wang, J., Raskin, L., Samuels, D. C., Shyr, Y., & Guo, Y. (2015). Genome measures used for quality control are dependent on gene function and ancestry. Bioinformatics (Oxford, England), 31(3), 318–323. doi: 10.1093/bioinformatics/btu668

Weider, L. J., Lampert, W., Wessels, M., Colbourne, J. K., & Limburg, P. (1997). Long-term genetic shifts in a microcrustacean egg bank associated with anthropogenic changes in the Lake Constance ecosystem. Proceedings of the Royal Society of London Series B-Biological Sciences, 264(1388), 1613–1618.

Wells, D., Sherlock, J. K., Handyside, A. H., & Delhanty, J. D. (1999). Detailed chromosomal and molecular genetic analysis of single cells by whole genome amplification and comparative genomic hybridisation. Nucleic Acids Research, 27(4), 1214–1218. doi: 10.1093/nar/27.4.1214

Wickham, H. (2016). ggplot2 Elegant Graphics for Data Analysis (Use R!). In Springer. doi: 10.1007/978-0-387-98141-3

Woyke, T., Sczyrba, A., Lee, J., Rinke, C., Tighe, D., Clingenpeel, S.,…Cheng, J.-F. (2011). Decontamination of MDA reagents for single cell whole genome amplification. PloS One, 6(10), e26161. doi: 10.1371/journal.pone.0026161

Xu, S., Ackerman, M. S., Long, H., Bright, L., Spitze, K., Ramsdell, J. S.,…Lynch, M. (2015). A male-specific genetic map of the microcrustacean *Daphnia pulex* based on single-sperm whole-genome sequencing. Genetics, 201(1), 31–38. doi: 10.1534/genetics.115.179028

Xu, Y., & Zhao, F. (2018). Single-cell metagenomics: challenges and applications. Protein & Cell, 9(5), 501–510. doi: 10.1007/s13238-018-0544-5

Zhang, F., Ding, Y., Zhu, C. D., Zhou, X., Orr, M. C., Scheu, S., & Luan, Y. X. (2019). Phylogenomics from low-coverage whole-genome sequencing. Methods in Ecology and Evolution. doi: 10.1111/2041-210X.13145

Zhang, L., Cui, X., Schmitt, K., Hubert, R., Navidi, W., & Arnheim, N. (1992). Whole genome amplification from a single cell: Implications for genetic analysis. Proceedings of the National Academy of Sciences of the United States of America. doi: 10.1073/pnas.89.13.5847

Zheng, X., Gogarten, S. M., Lawrence, M., Stilp, A., Conomos, M. P., Weir, B. S.,…Levine, D. (2017). SeqArray-a storage-efficient high-performance data format for WGS variant calls. Bioinformatics. doi: 10.1093/bioinformatics/btx145

Zheng, X., Levine, D., Shen, J., Gogarten, S. M., Laurie, C., & Weir, B. S. (2012). A high-performance computing toolset for relatedness and principal component analysis of SNP data. Bioinformatics, 28(24), 3326–3328. doi: 10.1093/bioinformatics/bts606

